# Effects of sampling methodology on phenology indices: insights from sites across India and modelling

**DOI:** 10.1101/2023.12.26.573352

**Authors:** Yadugiri Tiruvaimozhi, Karthik Teegalapalli, Abinand Reddy, Akhil Murali, Aparajita Datta, Aparna Krishnan, Jayashree Ratnam, Mahesh Sankaran, Shasank Ongole, Srinivasan Kasinathan, T. R. Shankar Raman, Geetha Ramaswami

## Abstract

Plant phenology is the study of timing and extent of leaf, flower, and fruit production. Phenology data are used to study drivers of cyclicity and seasonality of plant life-history stages, interactions with organisms such as pollinators, and effects of global change factors. Indices such as timing of phenological events, proportion of individuals in a particular phenophase, seasonality, and synchrony have often been used to summarise plant phenology data. However, these indices have specific utilities and limitations and may be sensitive to sampling methodology, making cross-site comparisons challenging, particularly when data collection methods vary in terms of sample size, observation frequency, and the resolution at which phenophase intensity scores/values are recorded. We use fruiting phenology data from tropical trees across five sites in India to study the effects of sampling methodology on two indices: an index of population-level synchrony (overlap), and an index of seasonality. We supplement these results with simulations of fast- and slow-changing phenologies to test for the effects of sampling methodology on these indices. We found that the overlap index is sensitive to the phenophase intensity measurement resolution—with coarser intensity measures leading to overestimation of the overlap index. The seasonality index, on the other hand, was not affected by intensity resolution. Simulations indicated that finer intensity resolution is more important than frequency of observation to accurately estimate population synchrony and seasonality for fast- and slow-changing phenophases. Based on our findings, we provide recommendations for study design of future tropical tree phenology research, particularly for long-term or cross-site studies.

## Introduction

Phenological research involves monitoring the timing of cyclic biological events which have ecological and evolutionary relevance (Augspurger 1983). It also involves understanding the proximate and ultimate causes of the observed phenological patterns. The timing of events such as leafing, flowering, and fruiting reflects the link between plant growth and reproduction and ecosystem characteristics including biotic variables such as pollination and dispersal, and abiotic variables such as temperature, rainfall, and other factors (Hemingway & Overdorff 1999). Insights from long-term plant phenological research can also enable the prediction of the effects of climatic changes on plants since climate change-driven phenological shifts can disrupt biotic interactions and thereby affect entire ecosystems (Fisogni et al. 2022).

Phenological synchrony, the overlap and similarity of a phenophase among individuals, and seasonality, for instance, the peak of a phenophase, have ecological and evolutionary relevance at various scales ranging from the individual to a community. Several indices of synchrony have been reported in the phenology literature, and these are typically calculated at the scale of an individual plant (e.g., Augspurger, 1983; Freitas & Bolmgren 2008; Bogdziewicz et al. 2020). A synchrony index may compare only the occurrence of a phenophase in an individual with respect to every other individual in a population (e.g., Augspurger index; Augspurger, 1983). It may also be a composite measure comparing both phenophase occurrence and intensity levels of an individual to those of all other individuals in the population (e.g., Freitas-Bolmgren index; Freitas & Bolmgren 2008). In the latter approach, maximum synchrony is achieved when an individual’s phenophase co-occurs with those of all other individuals in a population, and all individuals show the maximum phenophase (e.g., if all individuals are in full bloom). Synchrony declines as individuals in a population vary in their timing (and/or intensity) of a phenophase. Both aseasonal and seasonal populations can display high synchrony. Within a species, asynchronous ripening of fruits may reduce competition for dispersers or consumption by seed predators (McKey 1975; Wheelwright 1985; Terborgh 1990), whereas flowering synchrony may increase pollinator visitation and thereby seed set (Freitas & Bolmgren 2008; Bogdziewicz et al. 2020). Seasonality is particularly relevant in terms of the influence of climate change on phenological patterns which can further affect the global carbon cycle (Häninen & Tanino 2011).

The quantification of phenological synchrony and seasonality can therefore provide insights into potential underlying evolutionary mechanisms. Values of these indices can be expected to vary based on the sampling methods used, particularly sample size, frequency of observation, and the phenophase intensity scores/values (see Figure 1 for an illustration), and this can further influence the interpretation of phenological events of a species or populations. Nevertheless, only a handful studies have focused on the variation in phenological indices caused by differences in sampling methodology (e.g., Chapman et al. 1992, 1994, Freitas & Bolmgren 2008, Hemingway & Overdorff 1999, Morellato et al. 2010).

**Figure 1.**
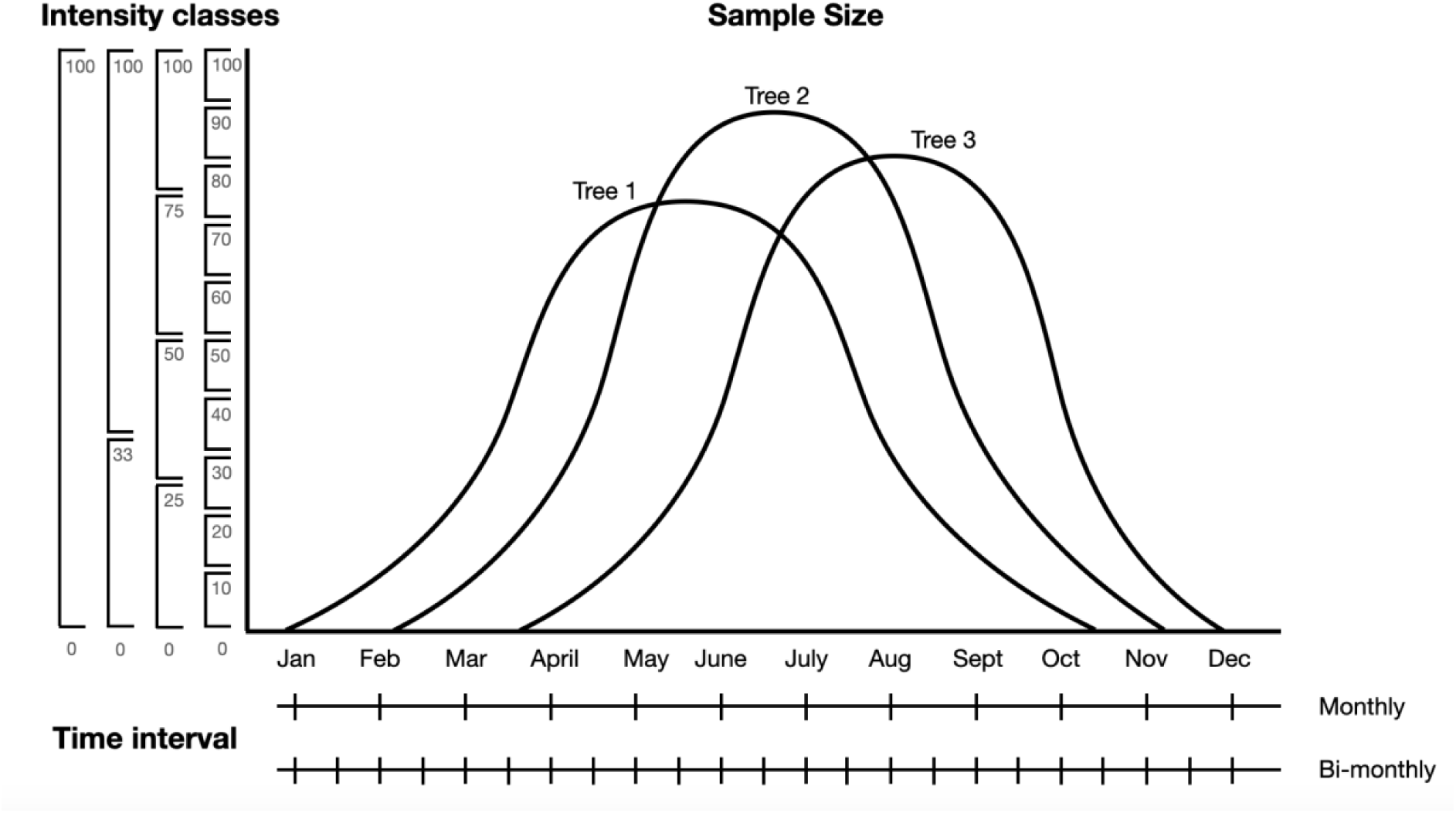
Variables related to sampling methods that affect phenology indices: resolution of intensity of phenology measurements (usually expressed as a percentage), sample sizes (number of individual trees observed per species), and time interval (frequency of phenology measurements).

We leveraged long-term phenological data on fruiting of trees and shrubs from field studies in four sites in India and from a citizen science effort for the entire state of Kerala (SeasonWatch Citizen Science Network, 2022) to study the effects of sampling methodology on select phenology indices. To supplement this, we also investigated the effects of variations in phenological methods on the indices using simulated data. In the simulations, the variations in the methods could be more robustly represented to arrive at implications for future phenological research. Our specific objectives were as follows:

1. What are the effects of differences in sampling methods on indices of phenological synchrony (overlap index) and seasonality?
2. What are the effects of differences in sampling methods on the indices of synchrony and seasonality, based on simulations of populations with varying rates of phenophase change (slow- and fast-changing phenophases)? For instance, in a population with a short-lived phenophase (fast-changing phenophase), phenological indices may be more sensitive to the frequency of observation, resolution of phenology intensity measurement, and/or sample size.

Based on the above results, we suggest optimal sampling parameters and methods to study tropical tree phenology across populations and communities.

## Materials and Method

### Study sites and datasets

We used fruiting phenology data (unripe and/or ripe fruits) from four tree phenology monitoring sites across India, and from a pan-India citizen science effort—SeasonWatch (Table 1). From SeasonWatch, data for the state of Kerala was chosen since it had the best spatial spread and representation of species as compared to other states of India. These sites differ in their climate with Pakke, Anamalais, and Kerala being wetter sites, and Rishi Valley (hereafter, RV), and Nagarjunasagar Srisailam Tiger Reserve (hereafter, NSTR) being drier sites. The sites also differ in seasonality and tree species composition (Table 1). Details about the habitat, species, effort, and methods are provided in Appendix 1.

**Table 1.** Description of the sites and datasets included in this study. NSTR: Nagarjunsagar Srisailam Tiger Reserve.

| Site | Date range | Number of species monitored | Sample size (no. individual trees per species) | Frequency of sampling | Resolution of phenology intensity measures | Elevation range (m) | Average annual rainfall (mm) |
| --- | --- | --- | --- | --- | --- | --- | --- |
| Rishi Valley | 2007–2016 | 18 | 10–40 | Fortnightly | 0, 33%,<br>100% (none, few, many) | ~700 | 690 |
| NSTR | 2018 | 44 | 10 | Fortnightly | 10% intervals | 500–850 | 713 |
| Pakke | 2011–2019 | 35 | 10–25 | Monthly | 25% intervals | 100–300 | 2500 |
| Anamalais | 2017–2020 | 172 | 1–41 | Monthly | 25% intervals | 800–1400 | 2500 |
| Kerala state (SeasonWatch) | 2014–2022 | 150 | variable | Weekly | 0, 33%,<br>100% (none, few, many) | 0–2500 | 3348 |

### Phenology methods

The methods used across the sites vary and here we focused on three key methodological properties: (a) Phenology intensity is a measure of the fraction of the tree crown showing a given phenology. The resolution of phenology intensity measurements varies, and can range from a presence/absence measure to values comprising 10% intervals in our datasets, (b) Sample size, which ranges from 2 to 41 individuals of a species in the site-based phenology datasets, to several thousand in SeasonWatch, and (c) Time interval or frequency of observations, which ranges from weekly to monthly (Table 1).

### Phenology indices

We focused on two frequently used indices in phenology studies to demonstrate the effects of methods used in phenology data collection on the final metric and consequently on potential inferences – the overlap index and seasonality index.

We consider an overlap index in this paper, which is intermediate between the Augspurger and Freitas-Bolmgren synchrony indices (Fisogni et al. 2022). Here, maximum synchrony is achieved when an individual’s phenophase co-occurs with that of every other individual at the same intensity, even if it is not the maximum intensity of the phenophase (e.g., if all individuals have 50% flowering at a given time). In effect, a population in which all individuals show a particular phenophase simultaneously at the exact same intensity (i.e., zero variation in the timing and intensity of a phenophase) has maximum synchrony. Average overlap synchrony across all the individuals in a sample provides an estimate of population-level overlap synchrony.

Specifically, the overlap index (O) of individuals *i*, *j* in a population of *S* individuals is:

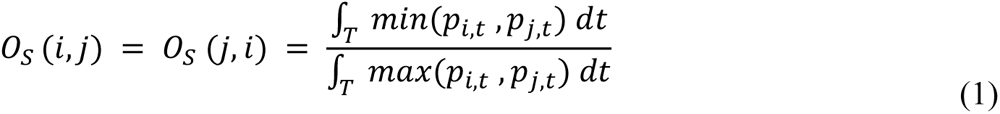

where, p = phenology measurement (e.g., intensity of fruiting), over time T.

Given that phenology measurements are taken at discrete time points in practice, the above formula can be written as follows, for time (t) increasing from 1 to T:

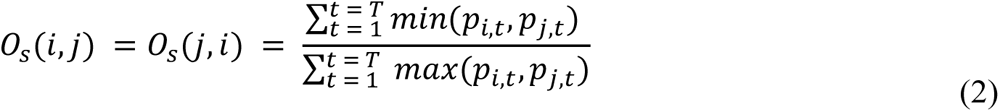

Overlap index of individual *i* in a population of *S* individuals is a composite value of the overlap of each individual *i* with every other individual in the population (or sample), and can be written as:

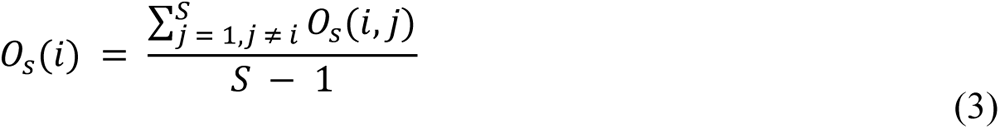

And population overlap is:

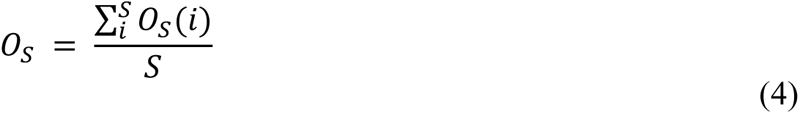

Seasonality of a phenophase for a given population is described by the average timing (day of the year) of the phenophase (which is the ‘peak’ of the given phenophase), and an index of amplitude, ρ, which describes how seasonal a given phenophase is, with larger ρ values indicating a highly seasonal phenophase. In this study, we refer to the index of amplitude, ρ, as the seasonality index. We use the function *circ.stats* in the R package CircStats to calculate both the timing and amplitude of seasonality for our datasets (Agostinelli 2018). Briefly, this package considers all the occurrences of a given phenophase in a year as vectors on a unit circle and calculates the mean direction (which is the average day of the year, or ‘peak’ season of the phenophase) and the mean vector length (which is the ‘seasonality index’ or ρ). Of these, we focus on the seasonality index, ρ, in this paper. The computation of both the seasonality index (ρ) and the overlap index used data on the occurrence of a phenophase, intensity of the phenophase, and sample size. In this paper, we assess the degree of influence of these factors on the computed indices.

### Simulated populations

In tropical forests of Southeast Asia and South America, variability in tree phenology has been found to result from cues such as extreme changes in solar radiation, rainfall, or temperature (Sakai et al. 1999, Butt et al. 2015). Accordingly, we chose temperature as a climatic trigger affecting reproductive phenology. To test the effects of methodological differences in data collection on phenology indices, we simulated two populations of 100 individuals each with a temperature-driven phenology, with temperature, and consequently the phenophase, peaking at about 180 days (Figure 2, a-b).

**Figure 2.**
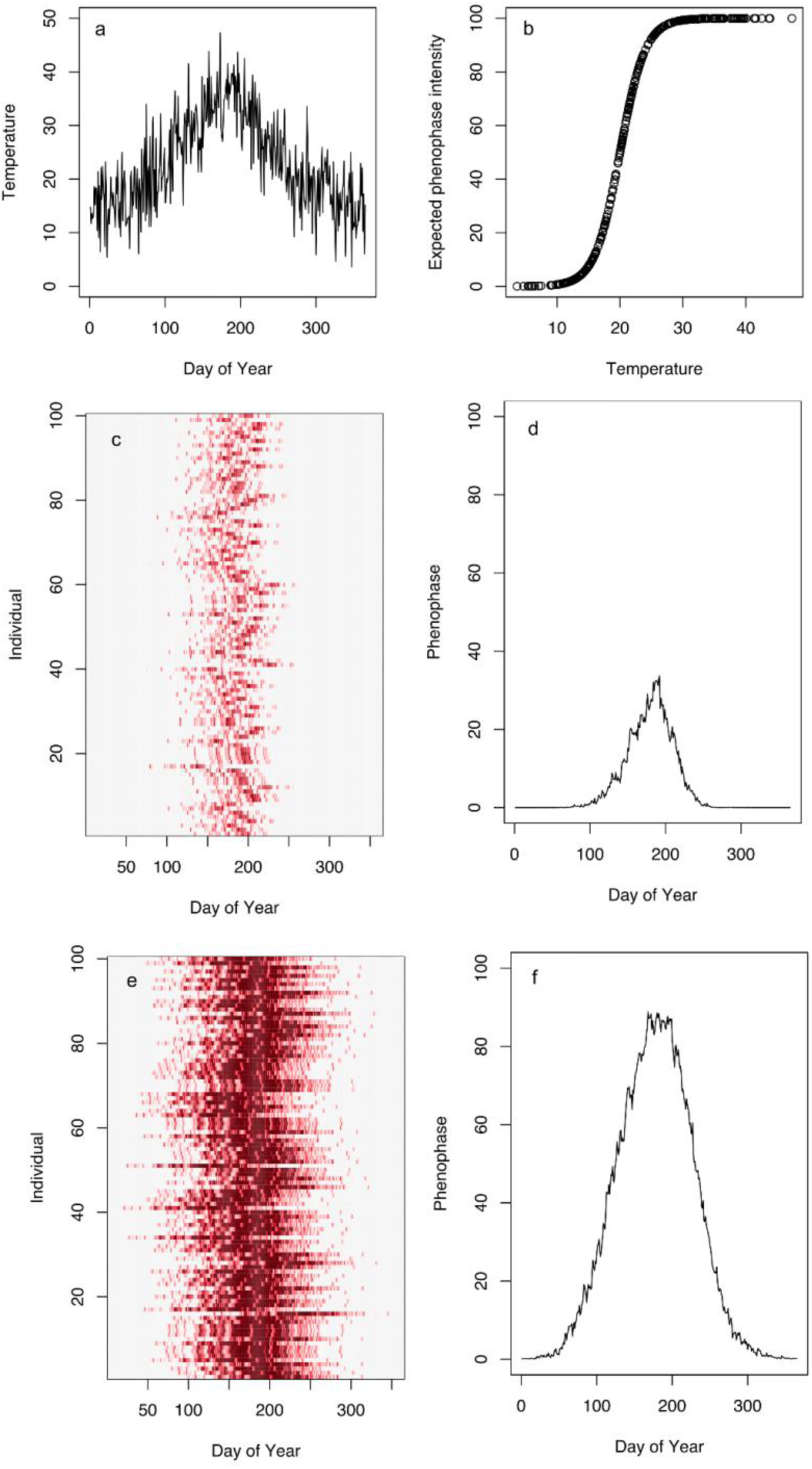
Simulation setup: a temperature-sensitive phenology was modelled, with temperatures peaking at 180 days (a, b). Phenologies of individuals and populations were simulated with a fast-changing phenophase (c, d) and a slow changing phenophase (e, f).

Temperatures were simulated to fluctuate from day to day by sampling from this distribution. The population phenologies respond to temperature with a lag of approximately 20 days. One of these populations was simulated to have a fast-changing phenophase, with individuals showing the phenophase for relatively fewer days. This was achieved by setting the sensitivity of the phenophase as 38°C. The second population was modelled to have a slow-changing phenophase, with temperatures above 28°C eliciting the phenophase (Figure 2, c-f).

### Analyses

For the primary datasets, we tested for the effect of phenology intensity resolution by comparing phenology indices calculated using actual data (fine-scale intensity resolution) with those calculated by setting intensity values greater than 0 to 1 (thus reducing it to a presence/absence measurement) (see Table 1 for details of phenological intensity resolution of the actual data from each site). In the simulated datasets, we compared the effect of recording phenology intensity at 10% intervals; 25% intervals; as none, few or many (0, 33, and 100%); and as absence/presence (0, 100%). We tested for sample size effects by iteratively sampling fewer individuals from the real and simulated datasets and using these data to compute phenology indices.

Overlap index across sites and species varied considerably. Most of the datasets did not represent the entire plant community, and since phenology was highly species-specific, we chose to examine representative species from these landscapes for further comparisons. To assess the effects of the resolution of phenological intensity on phenology indices, we chose the ten most observed species in each site, respectively (see Appendix 1 for details of representative species). To assess the effects of sample size in phenology data collection on phenology indices, we used phenology data for one representative species from each of the five datasets: *Erthroxylum monogynum* from Rishi Valley, *Anogeissus latifolia* from NSTR, *Horsfieldia kingii* from Pakke, *Paracroton pendulus* from Anamalais, and *Mangifera indica* from SeasonWatch.

## Results

### Phenology across sites – general trends

Reproductive phenology across sites, both in terms of overlap and seasonality indices, was site- and species-specific. In the two wetter sites (Pakke and Anamalais) and one seasonally-dry site (NSTR), for instance, fruiting in different species occurred almost throughout the year (Figure 3). In the two other datasets (RV and SeasonWatch), many of the observed species fruited around the same time of the year.

**Figure 3.**
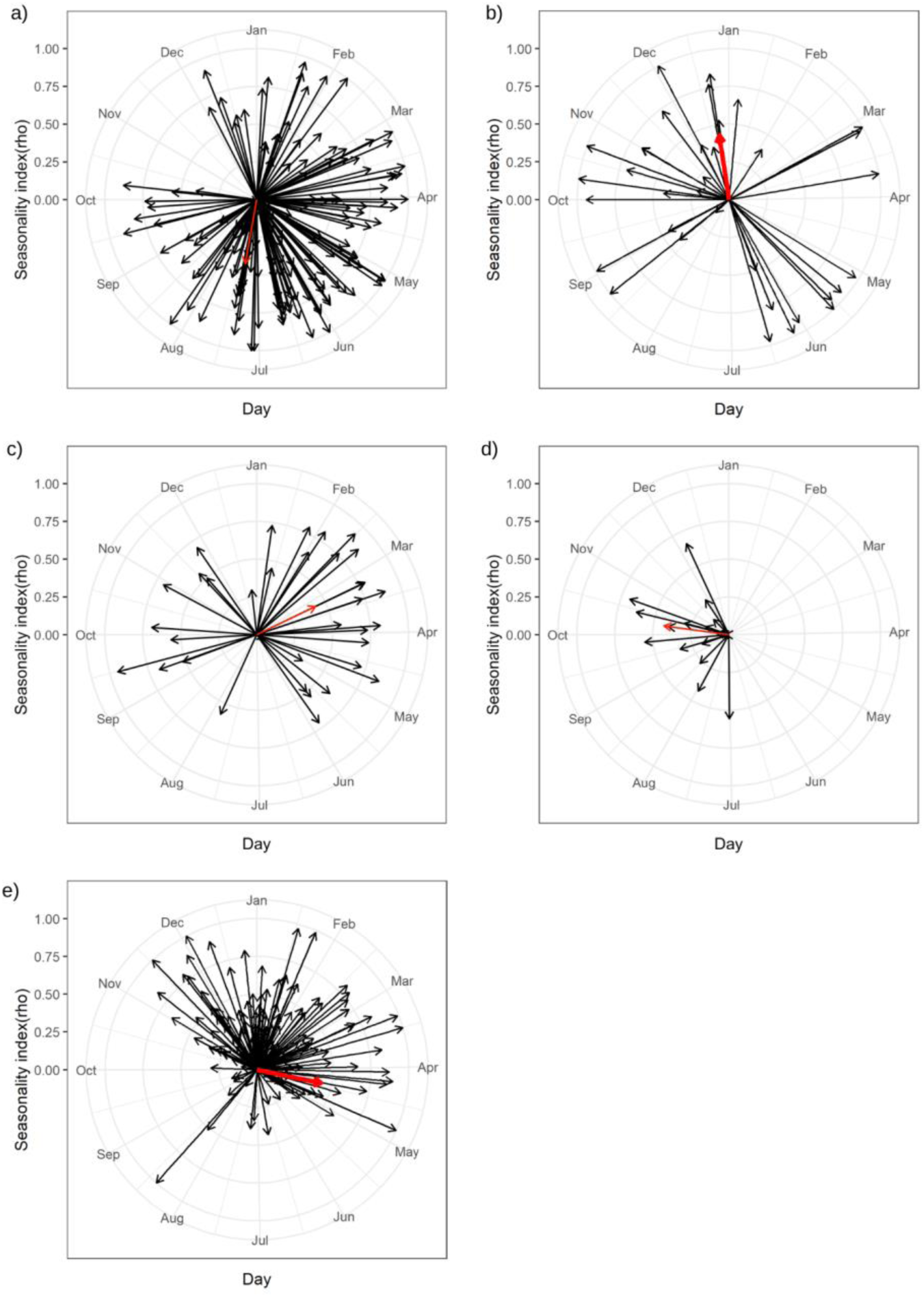
Seasonality (ρ) and timing of fruiting across five datasets from the tropics: a) Anamalais, b) NSTR, c) Pakke, d) RV, and e) SeasonWatch. Each arrow represents a species observed in the dataset, with the direction of the arrow indicating mean season of the phenophase, while the length of the arrow indicates the mean amplitude of the phenophase (longer arrow indicates more seasonal response). The red arrow represents seasonality values for a select species from each dataset – *Paracroton pendulus* (Anamalais), *Anogeissus latifolia* (NSTR), *Horsfieldia kingii* (Pakke), *Erthroxylum monogynum* (Rishi Valley), and *Mangifera indica* (SeasonWatch).

### Intensity resolution effects on phenology indices

Overlap index of fruiting in 10 representative species of Pakke and Rishi Valley varied considerably, from values close to zero overlap to high overlap (0.05–0.78 in Pakke, and 0.11–0.72 in Rishi Valley). Overlap indices in SeasonWatch and Anamalais datasets tended to be lower than 0.5 (0.01–0.2 in SeasonWatch, and 0.04–0.36 in Anamalais), while ranging from 0.02 to 0.53 in the NSTR dataset. In four out of five datasets, the phenological overlap index for fruiting was marginally higher when coarser intensity measurements were used for the 10 most observed species as compared to when finer intensity measures were used (Figure 4). The effect of resolution of intensity measurements on overlap was most apparent in the NSTR dataset, which had the finest resolution of intensity estimates for phenology (10% intervals). Here, the overlap index was substantially overestimated when intensity was measured at a coarse (presence/absence) scale.

**Figure 4.**
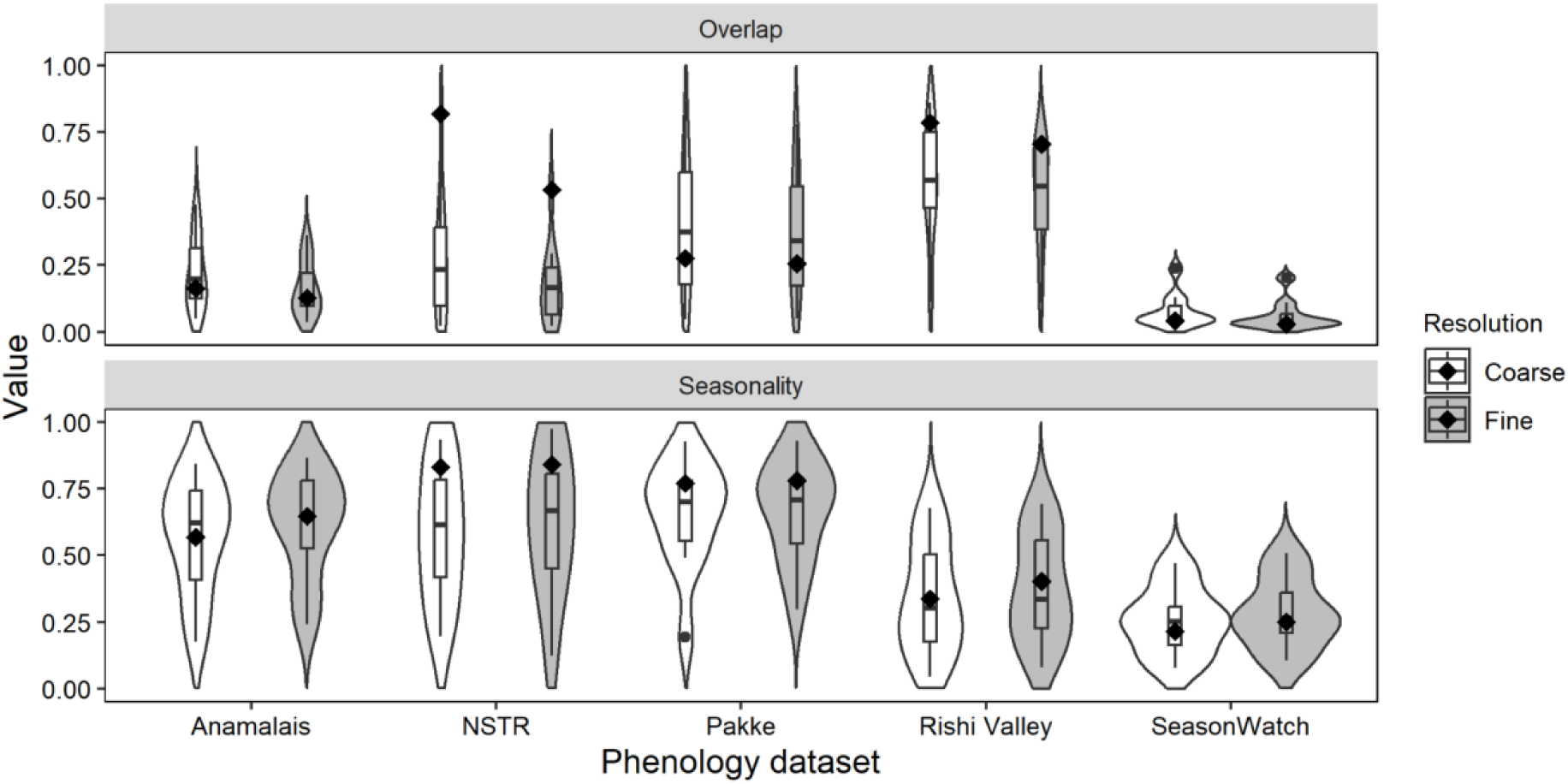
Overlap and seasonality index for fruiting in the 10 most observed or dominant species from five phenology datasets with two levels of phenology intensity resolution. The diamond within each violin plot indicates the mean overlap and seasonality index of one representative species from each site: *Paracroton pendulus* (Anamalais), *Anogeissus latifolia* (NSTR), *Horsfieldia kingii* (Pakke), *Erthroxylum monogynum* (Rishi Valley), and *Mangifera indica* (SeasonWatch). ‘Fine’ corresponds to the finest level of phenology intensity measurement i.e., 0, 10%, 20%…100% in NSTR; 0, 25%, 75%, 100% in Pakke and Anamalais; and 0, 33%, 100% in the Rishi Valley and SeasonWatch datasets, respectively. For all datasets, values above 0 were set to 1 in order to derive the ‘Coarse’ intensity measurement resolution, corresponding to a phenophase being present or absent. Error bars on the overlap index indicate standard errors across the population of the species observed in the dataset.

The seasonality index for 10 representative species varied considerably in four out of five datasets, but on average tended to be above 0.5, indicating that species were seasonal in their fruiting phenology. The SeasonWatch species on average had lower seasonality index because of the wider geographic scale of measurement. Seasonality index across datasets was not affected by resolution of phenology intensity measurements (Figure 4).

### Sample size effects on phenology indices

In the SeasonWatch dataset, in which the sample size of *M. indica* was 1824, sample size effects were strongly apparent; the overlap index was higher at low sample sizes as compared to mid- and high sample sizes (Figure 5). In all other datasets, the overall sample size for observation was between 5 and 41 trees and therefore sample-size effects were not as evident (Figure 5). Effects of sample size on seasonality index were also comparable across datasets with marginal variation between lower-, mid-, and maximum sample sizes in Pakke, Rishi Valley, and Anamalais. Lower sample sizes yielded considerably higher seasonality index values only in SeasonWatch data (Figure 5, Figure S1).

**Figure 5.**
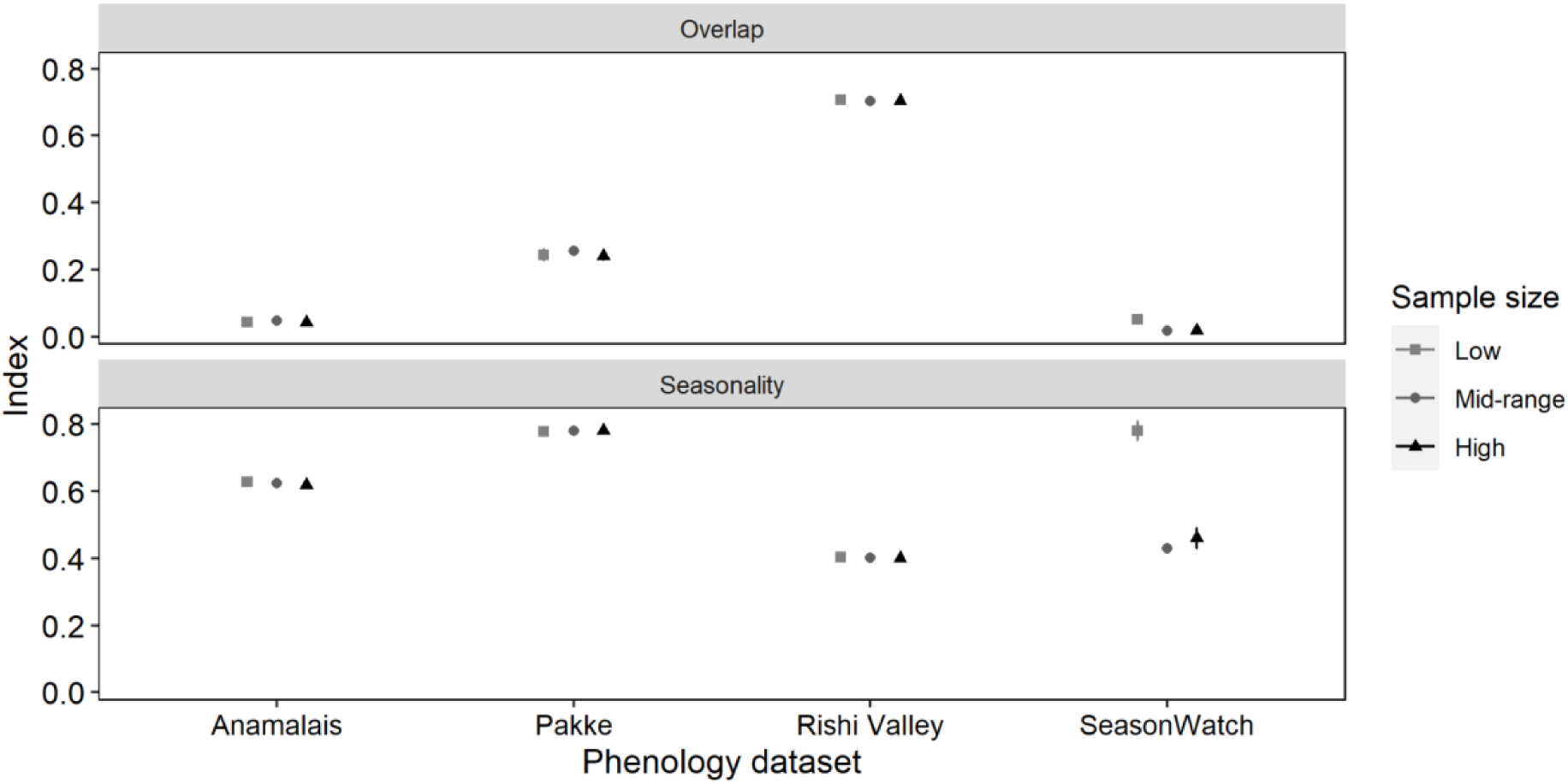
Effect of sample size (number of unique individuals monitored for phenology) on the overlap and seasonality index of fruiting in one representative species each across different phenology datasets. ‘Low’, ‘mid-range’, and ‘maximum’ in different datasets were as follows – Anamalais - 10, 20, 41; Pakke - 5, 10, 24; Rishi Valley - 10, 20, 40; and SeasonWatch - 30, 550, 1824. Error bars are based on 1000 random samples of low- and mid-range sample sizes. The effect of sample size was most discernible in the dataset with the highest maximum sample size (SeasonWatch) for overlap and seasonality, with both indices being over-estimated at lower sample sizes as compared to mid-range sample size. NSTR had overall species sample sizes of only 10 trees and was therefore excluded from this assessment.

### Effects of intensity resolution, sample size, and observation frequency in simulated populations

The simulated tree populations were of two types depending on the duration and amplitude of their phenophases. The simulated population with a fast-changing phenophase had short-duration, small-amplitude peaks (Figure 2c, d), and the population with a slow-changing phenophase had longer-duration, higher amplitude peaks (Figure 2e, f). Differences in phenology measurement methods affected overlap index estimates differently in these two populations. The overlap index for populations with a fast-changing phenophase was higher than the true value at low resolution measurements of phenophase intensity, but was relatively uninfluenced by frequency of observation or sample size (Figure 6a). The overlap index estimates were closest to the true value at the highest resolution of intensity measurements, irrespective of sample size or frequency of observation. The overlap index for populations with a slow-changing phenophase was also higher than the true value at low resolutions, but progressively approached the true value with increasing resolution of phenophase intensity measurements. Similarly, differences in sample size and frequency of observation showed no effect on estimates of overlap index. In both cases, greater resolution of phenophase intensity measurement captured the true overlap most accurately.

**Figure 6.**
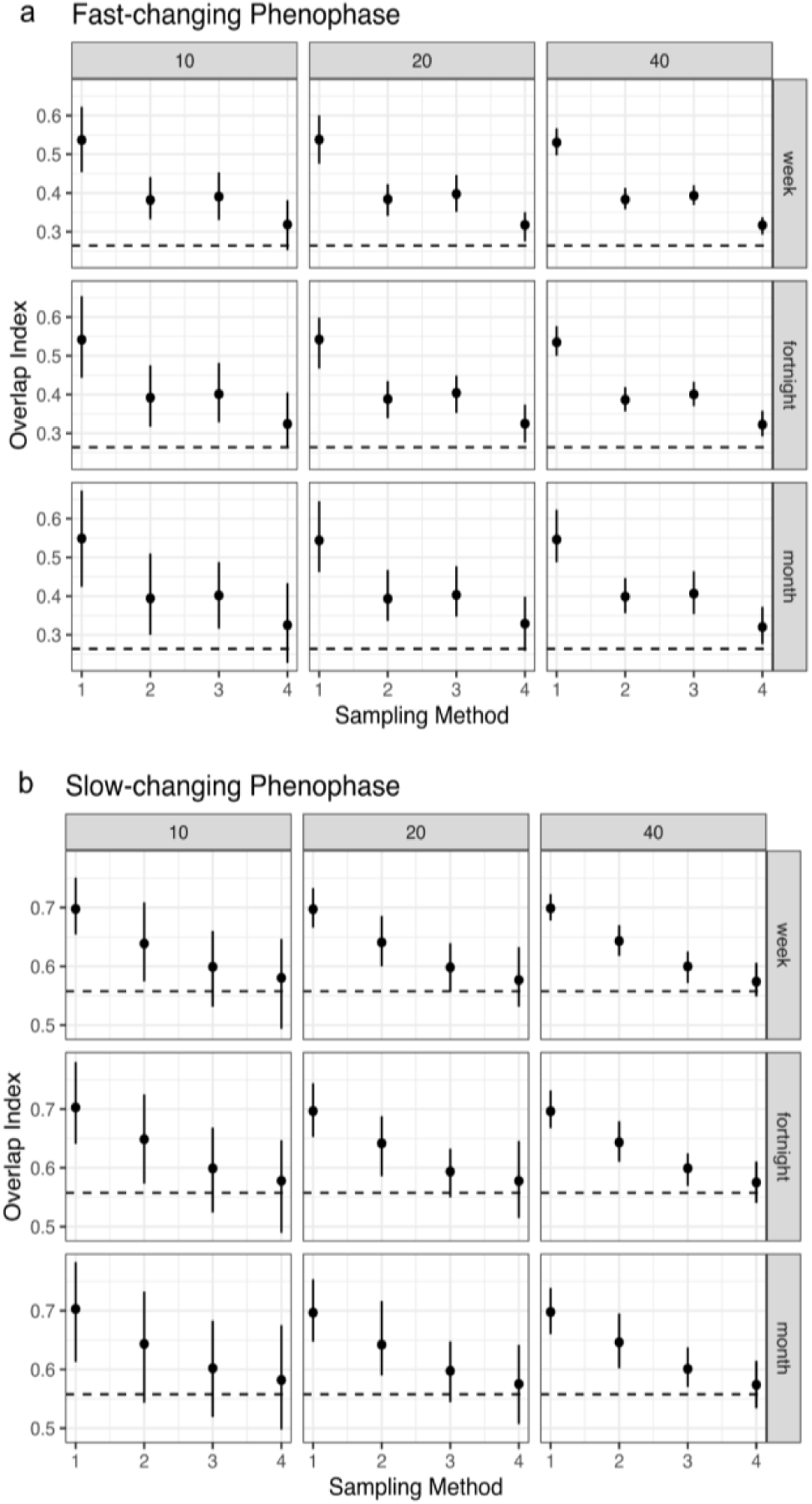
Sampling methodology effects on overlap synchrony index in simulated populations showing (a) fast-changing and (b) slow-changing phenology. Sampling methods (1, 2, 3, 4) correspond to phenology intensity measurements as follows: 1: presence/absence, 2: None, Few, Many, 3: 25% intervals, and 4: 10% intervals. Each column of panels corresponds to different sample sizes (10, 20, and 40 individuals), and rows represent frequency of sampling (i.e., whether the phenology data are recorded at weekly, fortnightly, or monthly intervals). The dashed horizontal lines represent the ‘true’ overlap synchrony of the two populations (0.264 for the population with the fast-changing phenology, and 0.557 for the slow-changing phenology population). Points are mean population-level overlap and error bars are confidence intervals calculated over 100 iterations.

As in the case of the overlap index, seasonality estimates were mainly affected by the resolution of phenophase intensity measurements (Figure 7a, b); they were underestimated at coarser intensity measures, and converged on the true value at finer-scale intensity measurements. This is in line with the results based on seasonality index calculations using site-based data as well (Figure 4). In the simulated data, frequency of observation and sample size do not appear to influence the estimated seasonality index for fast- and slow-changing phenologies. For both overlap and seasonality, larger sample sizes reduce the error around the average index estimates. Fast-changing phenologies have much larger errors around seasonality estimates than populations with slow-changing phenologies. This is mitigated to some extent at higher sample sizes. Overall, irrespective of whether observations were made at the weekly, fortnightly, or monthly scales, measuring intensity at a finer scale yielded better estimates of overlap and seasonality, whereas measuring more individuals yielded better precision of the estimates.

**Figure 7.**
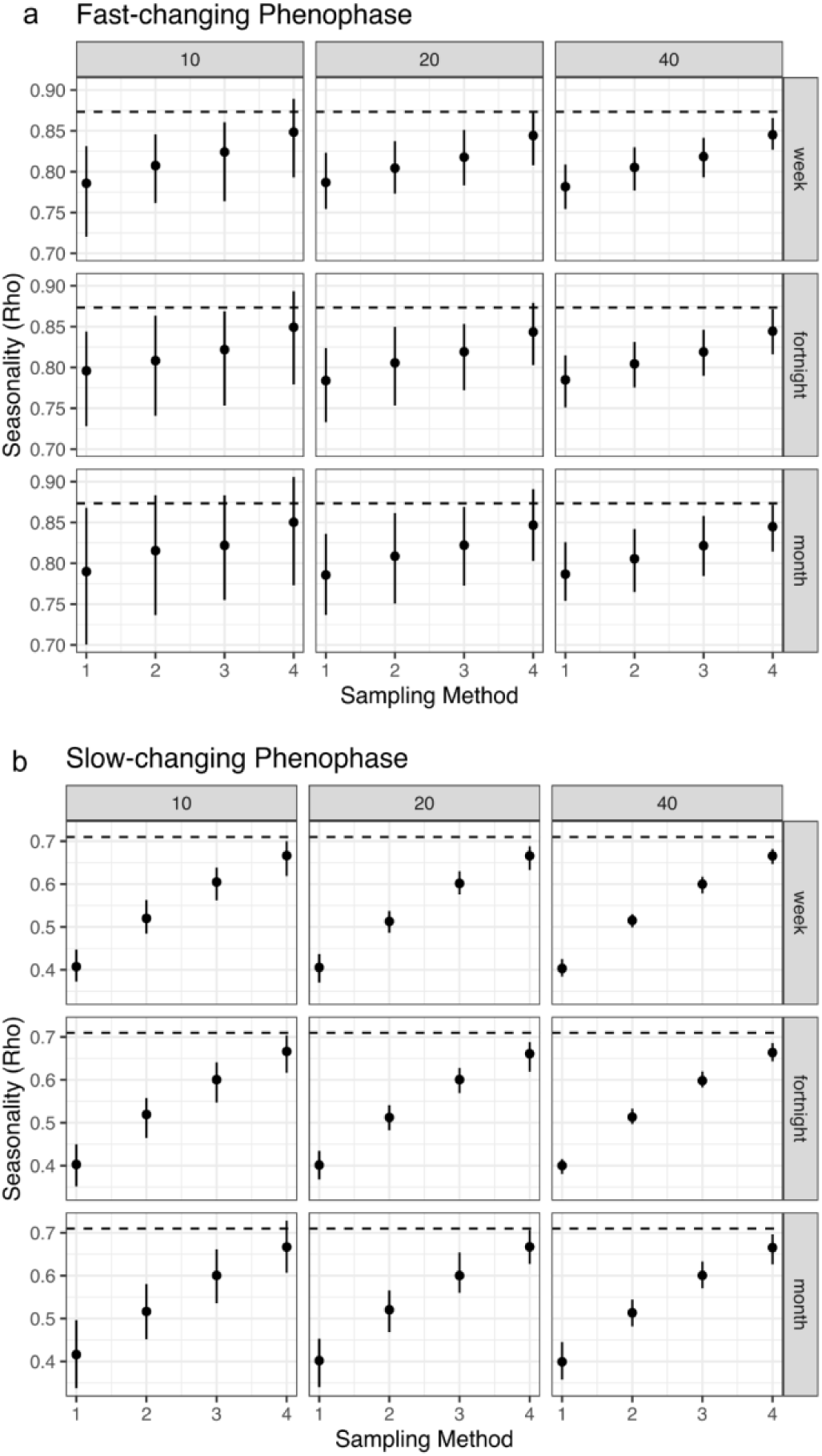
Sampling methodology effects on seasonality (ρ) index in simulated populations showing fast-changing (a) and slow-changing phenology (b). Sampling methods (1, 2, 3, 4) correspond to phenology intensity measurements as presence/absence; None, Few, Many; at 25% intervals; and at 10% intervals respectively. Each column of panels corresponds to different sample sizes (10, 20, and 40 individuals), and rows represent frequency of sampling (i.e., whether the phenology data are recorded at weekly, fortnightly, or monthly intervals). The dashed horizontal lines represent the ‘true’ ρ values of the two populations (0.873 for the population with the fast-changing phenology, and 0.710 for the slow-changing phenology population). Points are mean population-level seasonality and error bars are confidence intervals calculated over 100 iterations.

## Discussion

In this study, we sought to understand the effects of differing methodology on the estimation of two measures frequently used in plant phenological studies: overlap and seasonality indices. The primary motivation to conduct this study was to infer generalizable phenology trends across five datasets differing in methods of quantification, geographic location, and scale of phenological data collection. Overall, based on an analysis focused on a representative set of species per site, we found that only the finest level of intensity estimation (10% intervals) offered any advantage in the accurate estimation of the overlap index. Seasonality index across all datasets was not affected by the intensity resolution. As expected, depending on the overall variability in the population of interest, larger sample sizes of a single representative species allowed for higher precision in the estimation of overlap and seasonality across datasets. In simulated fast- and slow-changing populations as well, finer intensity resolution (10% intervals), and higher sample sizes (at least 40 individuals of each species) were required to accurately estimate these indices.

The NSTR dataset had the highest average overlap index in the fruiting phenology for representative species. This site is highly seasonal with a distinct wet period that lasts approximately four months with an average rainfall of 713 mm (1970–2015). Lobo et al. (2003) suggest that in forests with a distinct dry season, abiotic factors are possibly the proximate causes affecting phenological patterns, leading to a higher value of overlap. In Pakke and Anamalais, the overall higher moisture availability over a longer period possibly resulted in a wider seasonal window for fruiting, resulting in lower average overlap indices. Similar seasonal patterns of low overlap and wide distribution of fruiting across the year have been recorded in other tropical moist forest communities (e.g., Chapman et al. 2005; Wheelwright 1985).

Seasonality, on the other hand, was estimated as >0.4 in four out of five datasets, indicating that reproductive phenology in these species ‘peaked’ at certain times in the year. In the wetter sites, reproductive phenology was highly seasonal on average, while among the drier sites, NSTR was more seasonal than Rishi Valley. In both dry and wet sites, fruiting seasonality appeared related to the season of maximum rainfall, with the peak (as indicated by mean day of the year) occuring either just before (Pakke) or during (Anamalais) the main summer or south-west monsoon in the two wet sites.

We infer overlap and seasonality index values of the SeasonWatch dataset with more caution than other datasets. Owing to the large spatial spread, and inconsistent measurement of trees in the citizen science project, undocumented sources of variation in phenology overlap may make these indices inappropriate at this spatial scale. In a highly heterogeneous dataset such as SeasonWatch, it may be more reliable to use simple measures of phenology such as percent trees showing a phenophase across the spatial extent of the dataset. Other studies that have compared methodological effects on phenology concur that sampling strategies for site-specific studies need to be different from that of citizen science data (Morellato et al. 2018).

The simulation results indicate that standard indices of phenology can be affected by underlying species-specific phenology patterns. High variability in the timing and persistence of phenophases may be quantifiable using finest scales of intensity estimates, irrespective of frequency of observation and low sample sizes. Overlap and seasonality indices are both not ideal descriptors of phenology in rapidly changing, and short-duration phenology, except at the highest resolutions of phenology intensity measurements. An alternative to using absolute scales of intensity resolution, is to consider median values of intensity categories (e.g., 0, 0.16, 0.66 instead of 0, 0.33, 1). At median intensity resolution scales, all scales of intensity measurement other than presence/absence yield more accurate estimates of the true overlap (Figure S2).

We infer that an understanding of the underlying rate of phenophase change is essential to accurately estimate population-level synchrony in phenology using the overlap index. While studies elsewhere have highlighted the trade-off between sample size and frequency of observation (Morellato et al. 2010 and references therein), we highlight the role of underlying rate of change of a phenophase to be a key factor determining sampling methodology, irrespective of sample size and sampling duration (Figures 6, 7). This is particularly useful in the assessment of tropical tree species characterized by low sample size in forest communities (Morellato et al. 2010 and references therein). We found no effect of frequency of observation on the estimation of these two indices.

Both phenological seasonality and synchrony may be sensitive to temperature and precipitation changes and are therefore key to understanding plant responses to climate changes. Some plants may be more sensitive to climatic factors, and thereby have fast-changing phenophases. Other species may have a broader range of climatic conditions during which a phenophase is sustained, and thereby have slow-changing phenophases. We simulated these differences by manipulating the minimum temperature that stimulates a phenophase, with slower-changing phenophases stimulated at lower temperatures than fast-changing phenophases. We found that both slow- and fast-changing populations showed similar patterns in overlap and seasonality estimation when methods of phenological data collection differed. The clearest difference between slow- and fast-changing populations was that populations with fast-changing phenologies had much larger errors around seasonality estimates than populations with slow-changing phenologies. This suggested that larger sample sizes are more important in populations with fast-changing phenologies to accurately estimate seasonality. We expected frequency of observation to also become important in accurate overlap and seasonality estimation in populations with fast-changing phenophases, but did not find evidence for this in our simulations. Frequency of observation, however, may be an important factor in other phenology descriptors such as start or duration of phenophase, which are often used in cross-species or longitudinal phenology comparisons (Morellato et al. 2010).

We conclude that varying methodology and underlying phenology dynamics affect estimation of standard indices. Based on results from five diverse datasets and two simulated populations, we recommend the following best practices for analyzing phenological patterns using the overlap and seasonality indices:

a. For accurate estimation of the overlap index, finest intensity resolutions, at least at 10% intervals are needed: resolutions coarser than 10% intervals perform similarly to presence/absence data in the estimation of the overlap index.
b. Depending on the larger ecological question being addressed, seasonality may be a more appropriate index to characterise species phenology if fine-level intensity measurements are not logistically possible.
c. Pilot studies and examination of regional floras for approximate timing of phenophases can be used to determine whether the population of interest has fast- or slow-changing phenophases. In species with fast-changing phenophases, prioritizing higher sampling effort can yield more accurate estimates of the seasonality index.

## Acknowledgements

The authors would like to acknowledge the following people for collecting and collating data from different field sites: G. Moorthi, T. Sundarraj, T. Vanidas, A. Sathish Kumar, Kshama Bhat, P. Pavithra (Anamalais), Venkateshwarlu Byrapoghu, Niharika, and Ramesh (NSTR), Khem Thapa, late Kumar Thapa, late Tali Nabam, Narayan Mogar, Sagar Kino, Arjun Rai, Turuk Brah, Sital Dako, Amruta Rane, Ushma Shukla, Akanksha Rathore, Swati Sidhu, Rohit Naniwadekar, Veena Rai, Chaithra Gowda, Harman Kour, Noopur Borawake, Saniya Chaplod, Devathi Parashuram, Karishma Pradhan, Bibidishananda Basu (Pakke), P. Somnath (Rishi Valley), and the SeasonWatch Citizen Scientist Network comprising thousands of volunteers (SeasonWatch). The studies used in the paper were funded by Rohini Nilekani Philanthropies, AMM Murugappa Chettiar Research Centre (Anamalais), Department of Science and Technology GOI, National Centre for Biological Sciences, Bengaluru (NSTR), The Serenity Trust, Whitley Fund for Nature, National Geographic Society (Pakke), and the Wipro Foundation (SeasonWatch), as well as grants from Rainmatter Foundation, and philanthropic donations from Mr Arvind Datar. We wish to thank the Department of Environment, Forests & Climate Change in the states of Tamil Nadu, Arunachal Pradesh, and Andhra Pradesh and State Forest Departments for providing permissions and local support to conduct phenology studies.

---

**Figure S1.**
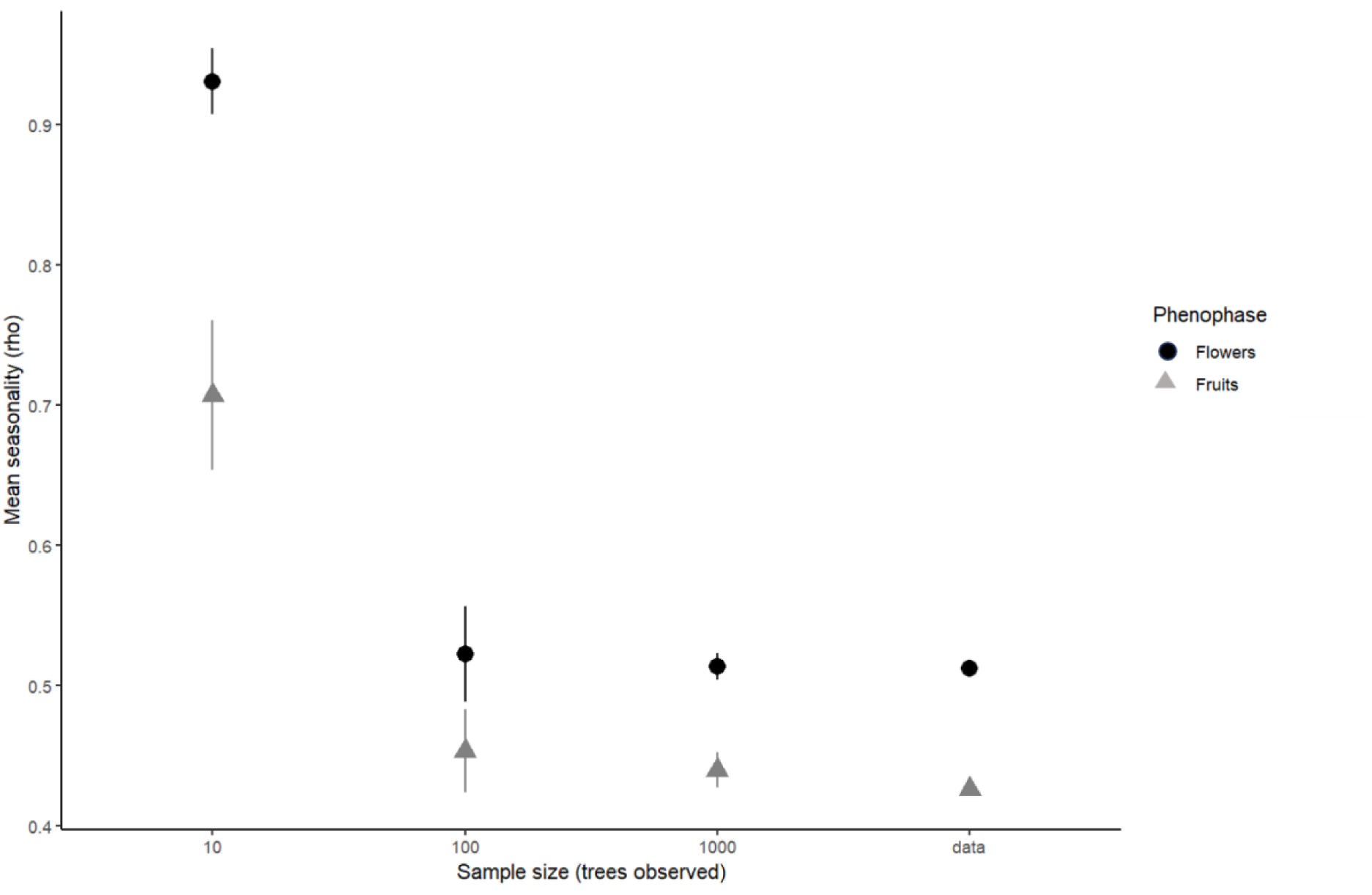
Sample size effects on seasonality index of fruiting in SeasonWatch data. *Mangifera indica* is one of the top three species observed on a weekly basis in Kerala, with a maximum sample size of 1824 trees in a year.

**Figure S2.**
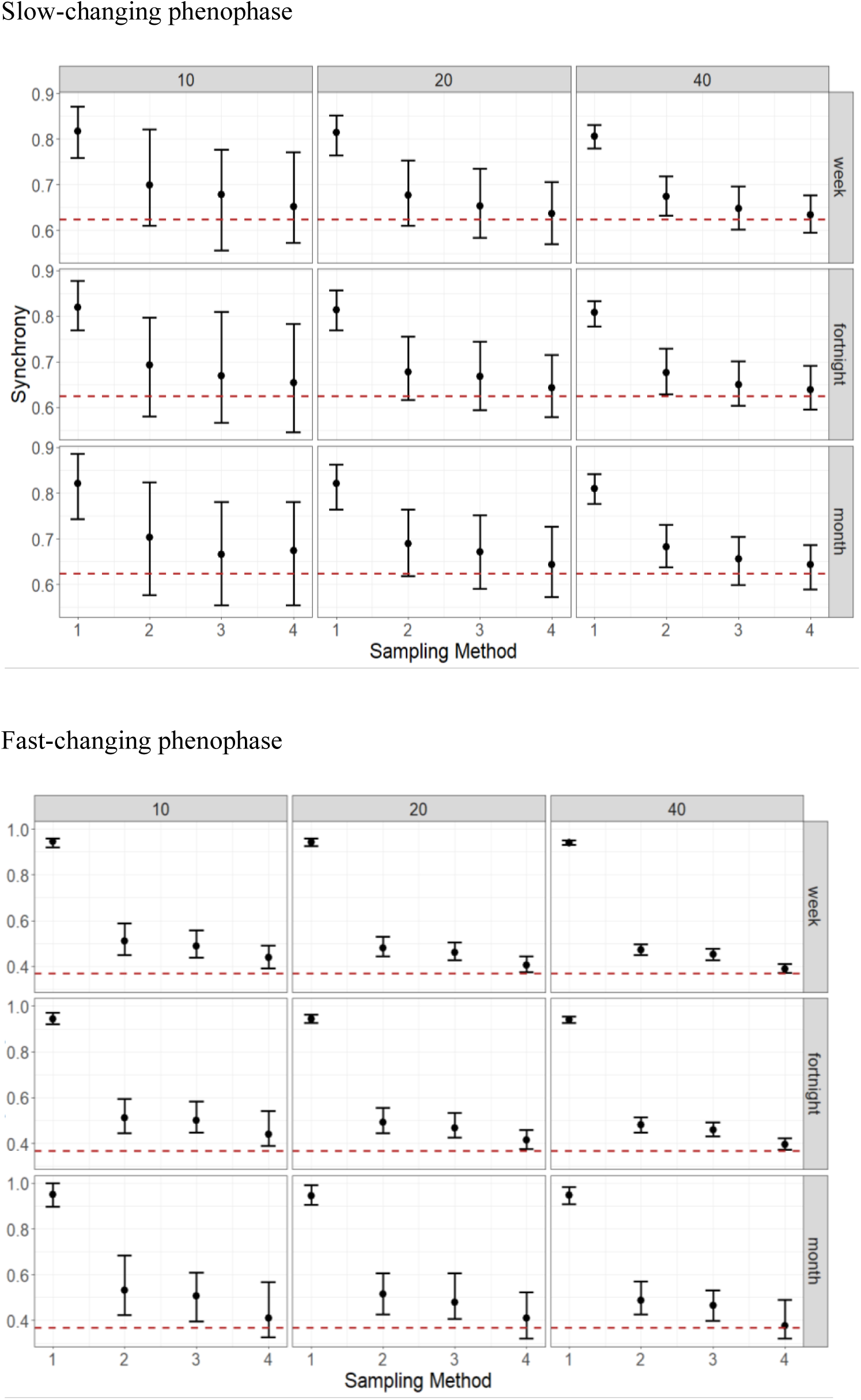
In simulated fast- and slow-changing phenophases, considering the median value of an intensity interval (e.g. 0, 0.16, and 0.66 instead of 0, 0.33, and 1) led to better estimates of population overlap index than exact intensity interval limits (in comparison with Figure 6).

## Appendix 1. Summary details of sampling effort and methods of phenology observation across the five datasets

### Pakke

A long-term tree phenology monitoring study of 722 reproductive trees of 53 selected species is underway in the tropical semi-evergreen forests of Pakke Wildlife Sanctuary and Tiger Reserve (see Datta & Rane 2013 for more detail regarding study site). The overall phenology tree sample represents 55% of the adult tree density and species composition at the study site based on prior studies (Datta 2001, Datta & Rawat 2008). Fifteen of the 20 top-ranked species are represented in the phenology sample. Other vegetation and climatic parameters are also recorded. Two to four observers carry out the monitoring every fortnight using binoculars to assess the presence of each phenophase in the tree canopies, along marked trails for each species. The phenophases recorded are the presence/absence of young leaf, mature leaf, senescent or old leaf, flower buds, open flowers, unripe fruit and ripe fruit. Unripe and ripe fruits are scored on a scale of 0 - 4, where 0 indicates no fruits and, 1 denotes 25% of canopy in fruit, 2 refers to 25 - 50% of the canopy in fruit, 3 indicates 50 - 75%, and 4 denotes 100% of canopy in fruit. For this paper, we used the monthly phenology data for 670 individual trees of 35 species (10 - 25 individuals per species) from April 2011 to September 2019. From this, data from 255 individuals of 12 species with 20 - 25 individuals per species were selected for representative species analyses. These representative species include *Aglaia* sp., *Ailanthus grandis, Dysoxylum cauliflorum, Dysoxylum gotadhora, Gynocardia odorata, Horsfieldia kingii, Livistona jenkinsiana, Pterospermum acerifolium, Sterculia villosa, Stereospermum chelonoides, Tetrameles nudiflora* and *Dalrympelea pomifera*.

### Anamalais

In total, 1376 individual trees of 172 species were observed monthly from March 2017 to December 2020. Phenological observations were made on 1 - 41 individuals per species along seven trails located on the Anamalais Plateau and Anamalai Tiger Reserve (10.3021°N, 76.8298°E – 10.4011°N, 76.9929°E) in the Anamalai Hills, Western Ghats. All trails were located in mid-elevation tropical wet evergreen forests of the *Cullenia exarillata* – *Mesua ferrea* – *Palaquium ellipticum* type (Pascal 1988). From this, data from 507 individuals of 13 species with 35–41 individuals per species were selected for representative species analyses. These representative species are *Acronychia pedunculata*, *Antidesma menasu*, *Cullenia exarillata*, *Euodia lunu-ankenda*, *Gomphandra coriacea*, *Litsea stocksii, Mesua ferrea*, *Myristica dactyloides*, *Palaquium ellipticum*, *Paracroton pendulus*, *Persea macrantha*, *Vateria indica*, and *Villebrunea integrifolia*. Trails were surveyed at the start of every month and trees were visually scored for the following phenophases: leaves (young/mature), flowers (buds, open), and fruits (unripe, ripe). Phenophase was scored on a scale of 0 to 4: 0 indicating absence of the phenophase, and values from 1 to 4 indicating the intensity (from low to high in 25% classes based on extent of canopy manifesting the phenophase).

### Rishi Valley

The study was carried out in a semi-arid tropical scrubland along the Eastern Ghats in the Deccan Plateau. The study area has distinct cool and warm, and dry and wet seasons (Ramaswami et al. 2019). Overall, 647 individual trees of 18 species (12–40 individuals per species) were observed from December 2007 to December 2016. From this, data from 600 individuals of 15 species with 40 individuals per species were selected for representative species analyses. These representative species are *Acacia leucophloea*, *Albizzia amara*, *Azadirachta indica*, *Chomelia asiatica*, *Delonix regia*, *Erythroxylon monogynum*, *Flacourtia sepiaria*, *Lantana camara*, *Peltoforum pterocarpum*, *Pongamia pinnata*, *Randia dumetorum*, *Santalum album*, *Strychnos nux-vomica*, *Tamarindus indica*, and *Wrightia tinctoria.* Trees in Rishi Valley are monitored every fortnight. Two stages of leaf (young, mature), two stages of flower (bud, open), and two stages of fruit (unripe, ripe) are monitored. The same observer has made these observations since 2007, and phenophase quantities are recorded as ‘none’, ‘few’, and ‘many’, corresponding to the phenophase being absent, detected in 30% of the canopy, or detected in >30% of the canopy.

### NSTR

The savanna woodland plot in NSTR is part of the long-term ecological monitoring network LEMoN India initiative (https://lemonindia.weebly.com). The site is located at an elevation of 700 a.s.l. and receives a long-term mean annual rainfall (1970–2015) of 713 mm, with eight dry months (Figure 1). Most of the rainfall occurs between July and October (summer monsoon). Overall, 113 individuals (girth at breast height > 10 cm) belonging to 11 species that comprised 80% of the stem basal area in a 1-ha plot were monitored fortnightly in the year 2018. The following variables were recorded: canopy fullness (L), and the percentage of flushing (FL), mature (M), and senescing leaves (S). During each session, we assigned a visual percentage score of 0 – 100 for each phenophase (flushing, mature, and senescence) with a resolution of 10%. The species are *Anogeissus latifolia*, *Bridelia retusa*, *Buchanania cochinchinensis*, *Chloroxylon swietenia*, *Dalbergia paniculata*, *Emblica officinalis*, *Eriolaena quinquelocularis*, *Grewia orbiculata*, *Pterocarpus marsupium*, *Terminalia elliptica*, and *Ziziphus xylopyrus*.

### SeasonWatch

This is a citizen science project collating information on tree phenology across India. In total, 120,000 individual trees of 170 species were observed either once or repeatedly every week from November 2011 to September 2023. From this, data from 47,224 individuals of 10 species with 1997–10238 individuals per species (1 Jan 2014–30 May 2022) were selected for representative species analyses. These species are: *Artocarpus heterophyllus*, *Mangifera indica*, *Tamarindus indica*, *Phyllanthus emblica*, *Cassia fistula*, *Tectona grandis*, *Syzygium cumini*, *Mimusops elengi*, *Bauhinia purpurea*, and *Samanea saman.* More than 85% of these data come from the south Indian state of Kerala (data used in this study). Trees are registered with the programme by citizen scientists and are observed on a weekly basis for three stages of leaf (young, mature, and dying), two stages of flower (bud, open), and two stages of fruit (unripe, ripe, and ‘open’ in dehiscent fruit species). Observers are trained to report phenophase quantities as ‘none’, ‘few’, and ‘many’, corresponding to the phenophase being absent, detected in 30% of the canopy, or detected in >30% of the canopy.

